# Personalized sleep-wake patterns aligned with circadian rhythm relieve daytime sleepiness

**DOI:** 10.1101/2021.03.15.435366

**Authors:** Jaehyoung Hong, Su Jung Choi, Se Ho Park, Hyukpyo Hong, Victoria Booth, Eun Yeon Joo, Jae Kyoung Kim

## Abstract

Shift workers and many other groups experience irregular sleep-wake patterns. This can induce excessive daytime sleepiness that decreases productivity and elevates the risk of accidents. However, the degree of daytime sleepiness is not correlated with standard sleep parameters like total sleep time, suggesting other factors are involved. Here, we analyze real-world sleep-wake patterns of shift workers measured by wearables with a newly developed user-friendly computational package that simulates homeostatic sleep pressure – the physiological need for sleep – and the circadian rhythm. This reveals that shift workers who align sleep-wake patterns with their circadian rhythm have lower daytime sleepiness, even if they sleep less. The alignment, quantified by a new parameter, circadian sleep sufficiency, can be increased by dynamically adjusting daily sleep durations according to varying bedtimes. Our computational package provides flexible and personalized real-time sleep-wake patterns for individuals to reduce their daytime sleepiness and could be used with wearable devices to develop smart alarms.

## Introduction

In our modern 24-h society, approximately 20% of the working population is engaged in shift work but more than 80% of the population has a shift work-like lifestyle with atypical light exposure (Sulli et al., 2018). Irregular sleep-wake patterns cause shift work disorder with symptoms including fatigue, sleepiness, insomnia, and poorer mental agility (Drake et al., 2004; Foster, 2020). In particular, excessive daytime sleepiness (EDS) reduces performance efficiency, increases the risk of work-related injuries, and is a significant public health problem (James et al., 2017; Slater and Steier, 2012). One way to reduce EDS could be to increase sleep duration. However, significant correlations between sleep durations or the other standard sleep parameters, including sleep efficiency and sleep latency, with daytime sleepiness of shift workers have not been identified (Gumenyuk et al., 2015; Kato et al., 2012). Furthermore, there have also been no connections identified between EDS and broader clinical features or features measured by polysomnography, i.e., in-depth sleep studies (Eiseman et al., 2012). This suggests the involvement of other, unknown factors in mediating the effects of irregular sleep-wake patterns.

The effect of irregular sleep-wake patterns on sleepiness has also been investigated with mathematical models (Abel et al., 2020; Van Dongen, 2004). While the details between the models differ, they are mainly based on the two-process model (Borbély, 1982), which simulates sleep-wake patterns according to interactions between homeostatic sleep pressure (the physiological need for sleep, which appears to be mainly determined by the level of somnogens such as cytokines, prostaglandin D2 (PGD2), and adenosine (Shi and Ueda, 2018; Skeldon et al., 2017)) and the circadian (~24 h) rhythm of the master clock in the suprachiasmatic nucleus. By simulating homeostatic sleep pressure and the circadian rhythm, the models successfully predicted subjective sleepiness and fatigue measured during long-lasting sleep deprivation in laboratory studies (Daan et al., 1984; Postnova et al., 2018; Puckeridge et al., 2011; Van Dongen, 2004), and irregular real-world work schedules (Moore-Ede et al., 2004; Van Dongen, 2004). While these model predictions suggested work schedules avoiding strenuous activities during times of high sleepiness to improve performance and minimize risks (Moore-Ede et al., 2004; Postnova et al., 2014), their widespread application is challenging without continual reinforcement (i.e., forcing a specific schedule) (Czeisler et al., 1982). Importantly, shift workers even with similar work schedules have dramatically different sleep-wake patterns and thus different daytime sleepiness (Vetter et al., 2015). This demonstrates a need for personalized and flexible sleep-wake schedules to prevent EDS.

Here, we analyzed the relationship between daytime sleepiness and real-world sleep-wake patterns of individual shift workers measured by wearable wrist actigraphy. Specifically, to analyze the complex individual sleep-wake patterns by simulating underlying homeostatic sleep pressure and the circadian rhythm, we developed a publicly accessible user-friendly computational package based on a physiologically-based mathematical model of sleep-wake cycles (Phillips et al., 2010; Skeldon et al., 2017; Swaminathan et al., 2017). This analysis revealed that as sleep-wake patterns became more aligned with an individual’s circadian rhythm, daytime sleepiness decreased, even if total sleep times were similar. To effectively investigate this relationship, we developed a new sleep parameter that we call Circadian Sleep Sufficiency (CSS). CSS was strongly correlated with daytime sleepiness, unlike other standard sleep parameters. CSS can be increased by adaptively adjusting daily sleep duration according to the personal choice of bedtime day-by-day rather than by forcing a specific work and sleep-wake pattern, thus providing a flexible and personalized solution to reduce daytime sleepiness. The personalized sleep-wake patterns can be provided in real-time when the user-friendly computational package developed in this study is linked with wearable devices.

## Results

### Daytime sleepiness is not significantly correlated with standard sleep parameters

We measured the activity and light exposure of 21 rotating nurses from Samsung Medical Center (SMC) every 2-min over 13 days using wrist activity monitors (Figure 1A and Table S1). Then, in each 2-min epoch, the status of the participants was categorized as either wake and active, wake and rest, sleep and active, or sleep and rest with Actiware-Sleep software whose accuracy has been validated previously (Edinger et al., 2004; Kushida et al., 2001). This allowed us to calculate six major standard sleep parameters: time in bed (TIB), sleep latency (SL), total sleep time (TST), wakefulness after sleep onset (WASO), number of awakenings (#Awak) and sleep efficiency (SE). We expected that as daily sleep duration (i.e., TST) increased, shift workers would be getting as much sleep as they needed, and this would decrease their daytime sleepiness, which was measured by the Epworth sleepiness scale (ESS). However, TST was not significantly correlated with daytime sleepiness (*P* = 0.50; Figure 1B). In particular, the daytime sleepiness of shift workers who had similar TST (6-7 h) differed dramatically. The other sleep parameters were also not significantly correlated with daytime sleepiness (Figures 1C, 1D, and S1A-S1D). The partial correlation between the standard sleep parameters and daytime sleepiness controlling for demographics of nurses (e.g., Age, BMI and the number of shift schedules) is also not significant (Table S1 and Figure S1E). Similarly, a previous study has also reported that the standard sleep parameters may not have a strong correlation with the daytime sleepiness of shift workers (Kato et al., 2012). This indicates the need for a different approach to analyze dynamic and complex sleep-wake patterns of shift workers.

**Figure 1.**
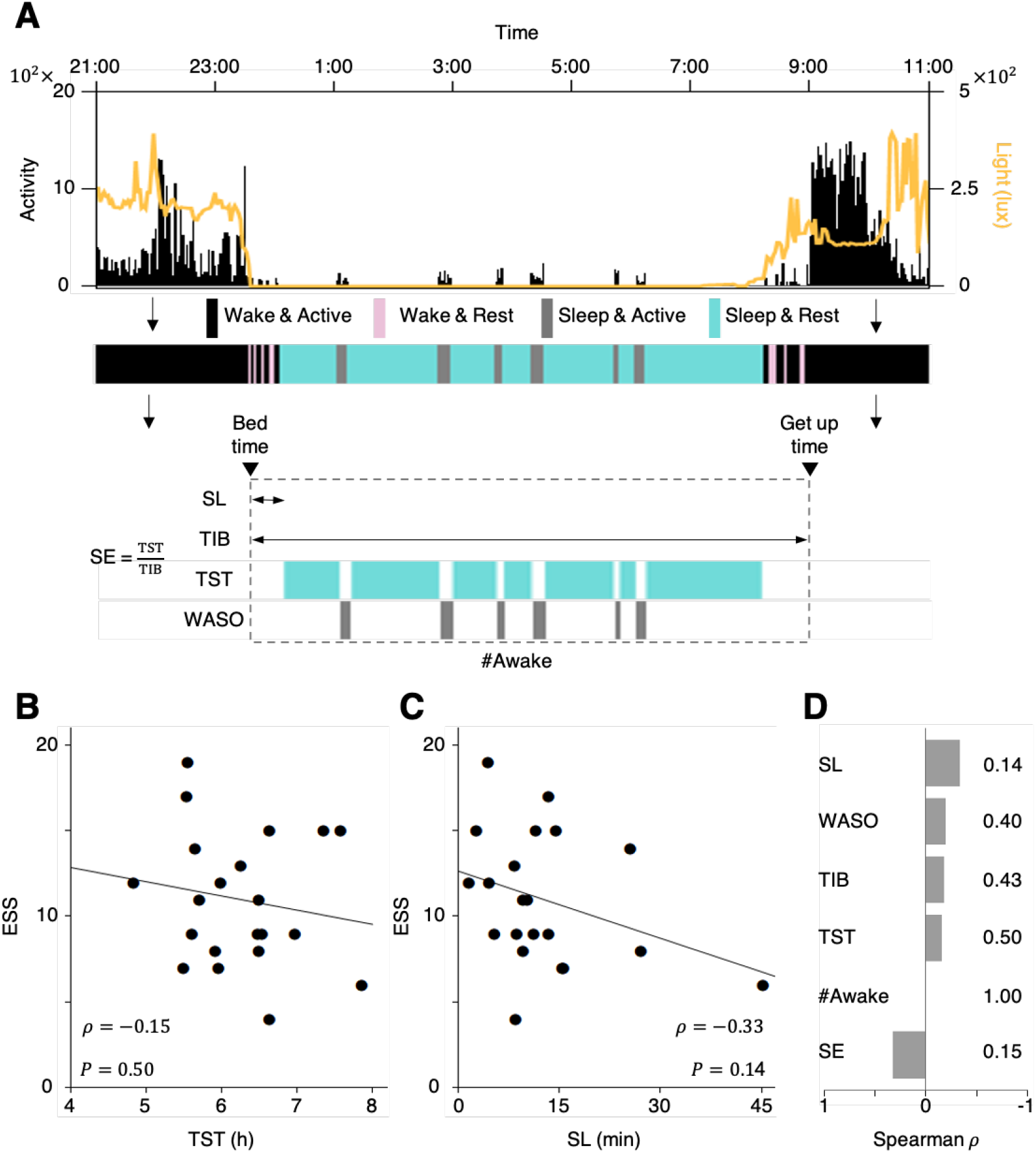
No significant correlation between standard sleep parameters and daytime sleepiness of shift workers. **(A)** Using activity (black vertical lines) and light exposure (yellow line) measured by the wrist actigraphy, the status of participants over time was categorized as either wake and active (black shade), wake and rest (pink shade), sleep and active (gray shade), or sleep and rest (blue shade) with Actiware-Sleep software, and then the six standard sleep parameters were calculated. TIB: time in bed; SL: sleep latency; TST: total sleep time; WASO: wakefulness after sleep onset; #Awak: number of awakenings; SE: sleep efficiency. **(B** and **C)** Scatter plots of TST (B) and SL (C) versus ESS of shift workers (*n* = 21). See Figure S1A-S1D for scatter plots of the other sleep parameters. Shift workers with similar TST (e.g. 6-7 h; B) had dramatically different daytime sleepiness. The line represents the least-square fitting line. *ρ* and *P* denote the Spearman’s rank correlation coefficient and *p* value of Spearman’s rank correlation test, respectively. **(D)** Correlations between the six standard sleep parameters and daytime sleepiness of shift workers were weak and not significant.

### A mathematical model is adopted to analyze dynamic sleep-wake patterns

To analyze the sleep-wake patterns of shift workers systematically, we modified a physiologically-based mathematical model of human sleep-wake cycles (Phillips et al., 2010; Skeldon et al., 2017; Swaminathan et al., 2017) (see Transparent Methods). In the model, the activity of sleep- and wake-promoting neurons, and thus sleep timing and duration, are determined by the interaction between the homeostatic sleep pressure and the circadian rhythm (Figures S2 and S3). The homeostatic sleep pressure describes the physiological need for sleep, which increases during wake and dissipates during sleep (black line; Figure 2A). The circadian rhythm, entrained to the external day-night cycle, determines the sleep and wake thresholds (yellow lines; Figure 2A). When the homeostatic sleep pressure increases above the circadian sleep threshold, the transition from wake to sleep is triggered (Figure 2A (i)). Thus, the model would naturally fall asleep whenever the homeostatic sleep pressure is higher than the circadian sleep threshold (i.e., in the gray ‘sleep region’; Figure 2A). To simulate wakefulness in the sleep region, similar to when shift workers work through the night, we modified the model to incorporate a ‘forced wakefulness’ (Figure 2A (ii) and Figure S4) (Phillips and Robinson, 2008; Postnova et al., 2014). When the homeostatic sleep pressure passes below the circadian sleep threshold (i.e., in the white ‘potential wake region’; Figure 2A), the transition from sleep to wake can occur naturally without forced wakefulness (Figure 2A (iii) and Figure S5). When the homeostatic sleep pressure further decreases below the circadian wake threshold (Figure 2A (iv)), the transition from sleep to wake is actively triggered. To simulate sleep (e.g. oversleeping or nap) in this case when the model would be naturally awake, we modified the model to incorporate a ‘forced sleep’ (Figure 2A (v) and Figure S4) (Phillips and Robinson, 2008; Postnova et al., 2014). Note that due to the lower level of the circadian sleep threshold during the night compared to the day (Figure 2A), falling asleep during the night occurs at a lower level of homeostatic sleep pressure (i.e., easier to sleep).

**Figure 2.**
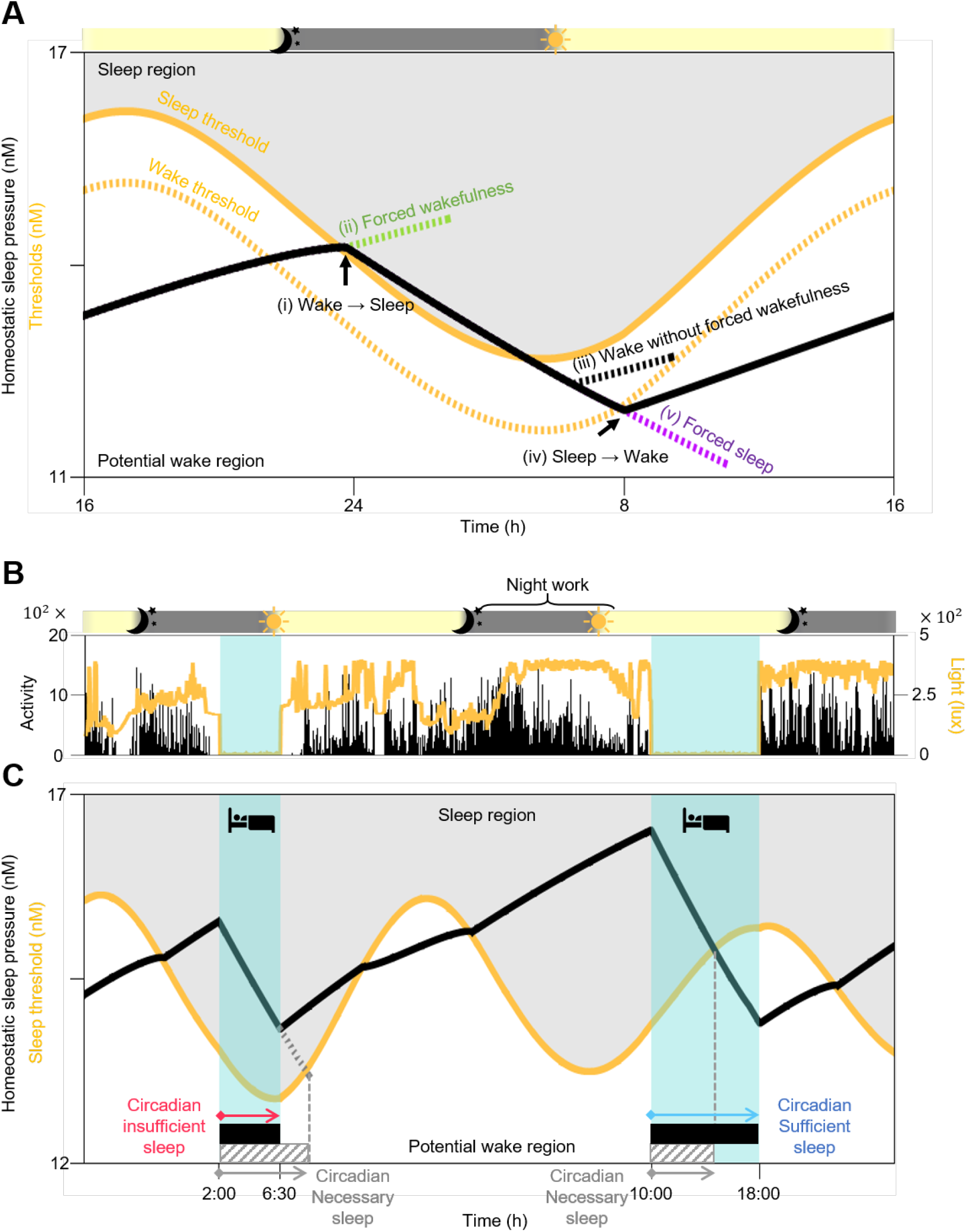
Sleep episodes are categorized as either circadian sufficient or circadian insufficient with a physiologically-based mathematical model of sleep-wake cycles. **(A)** In the mathematical model, the homeostatic sleep pressure (black line) dissipates during sleep and increases during wake. When it becomes higher than the circadian sleep threshold (yellow solid line), a transition from wake to sleep occurs (i). When the model would naturally fall asleep in the sleep region (gray region), forced wakefulness is needed to simulate wakefulness (ii). On the other hand, when homeostatic sleep pressure falls below the circadian sleep threshold and thus the model is in the potential wake region (white region), wakefulness can be simulated without forced wakefulness (iii). When the homeostatic sleep pressure falls below the circadian wake threshold (yellow dotted line), a transition from sleep to wake actively occurs (iv). In this case, forced sleep is required to simulate sleep (v). See Figures S2-S5 for details. Gray and yellow shades on top indicate the night (22:00-6:00 h) and the day (6:00-22:00 h), respectively. **(B** and **C)** The computational package based on the mathematical model simulated homeostatic sleep pressure (black line; C) according to sleep-wake patterns (blue shade; B), which were estimated by measured activity (black vertical lines; B). It also simulated the circadian variation of the sleep threshold (yellow line; C) by estimating the light signal reaching the circadian clock based on measured light exposure (yellow line; B). Then, the minimum sleep duration required to wake-up specifically in the potential wake region (i.e., the circadian necessary sleep; gray striped bars; C) was calculated for each sleep episode. Compared to circadian necessary sleep, longer or shorter sleep episodes (black bars; C) are referred to as circadian sufficient or circadian insufficient sleep, respectively. Gray and yellow shades on top of (B) indicate the night (22:00-6:00 h) and the day (6:00-22:00 h), respectively.

### A new sleep parameter, circadian sleep sufficiency, is strongly correlated with daytime sleepiness

With the modified mathematical model, we developed a publicly accessible user-friendly computational package (Figure S6) that simulates an individual’s homeostatic sleep pressure based on real-world sleep-wake patterns (blue shade; Figure 2B) that were mainly estimated by the wrist activity monitor (see Transparent Methods). Specifically, for the individual illustrated in Figure 2, the simulated homeostatic sleep pressure increased and decreased during wake and sleep, respectively, as expected (black line; Figure 2C). In particular, the homeostatic sleep pressure became extremely high after night shift work before the second sleep episode. Furthermore, the computational package estimated the light signal transmitted to the circadian clock based on the measured light exposures of the participant over time (yellow line; Figure 2B), and then simulated the circadian variation of the sleep threshold entrained to these light-dark cycles (yellow line; Figure 2C). The overall level of the simulated sleep threshold increased when exposed to light during the day or during the night shift, which inhibits falling asleep.

Based on the homeostatic sleep pressure and the sleep threshold simulated with the computational package, the duration of necessary sleep needed to wake up without effort can be predicted. Specifically, when people fall asleep, the computational package predicted how long they need to sleep so that their homeostatic sleep pressure decreased below their sleep threshold (into the potential wake region; Figure 2C) and thus they could wake up without effort (i.e., without forced wakefulness). We defined “circadian necessary sleep” as the sleep episode with the minimum duration required so that awakening occurs in the potential wake region (gray striped bars; Figure 2C). The duration of circadian necessary sleep depends on when individuals fall asleep and the subsequent intersection between their homeostatic sleep pressure and their sleep threshold, which are linked with their circadian rhythm. In the example in Figure 2C, the duration of the first sleep episode (black bar) is shorter than the duration of the predicted circadian necessary sleep (gray striped bar). This represents a situation when the individual wakes up before his/her homeostatic sleep pressure falls below the sleep threshold – i.e., forced wakefulness. We refer to this sleep episode as “circadian insufficient sleep” throughout the study. In contrast, the duration of the second sleep episode is longer than the duration of the predicted circadian necessary sleep for that cycle, meaning that the individual awakens several hours after their need for sleep has fallen below their sleep threshold. This is referred to as “circadian sufficient sleep” throughout the study.

We hypothesized that having circadian insufficient sleep – i.e., sleep accompanied by forced wakefulness–causes EDS. To investigate this, we compared the sleep-wake patterns of three shift workers who had considerably different daytime sleepiness measured by ESS despite having similar TST (6.65-6.98 h). Specifically, we categorized their daily sleep episodes (black bars; Figure 3A) as either circadian sufficient sleep (blue shade; Figure 3A) or circadian insufficient sleep (pink shade; Figure 3A) after comparing with their predicted circadian necessary sleep (gray striped bars; Figure 3A). The fractions of circadian sufficient sleeps in total sleep episodes were dramatically different between the three shift workers (from 81.25 to 69.23 to 50%) although they all had similar TST. Notably, as the fraction of circadian sufficient sleeps decreased, daytime sleepiness measured by ESS increased from 4 to 9 to 15 (Figure 3A).

**Figure 3:**
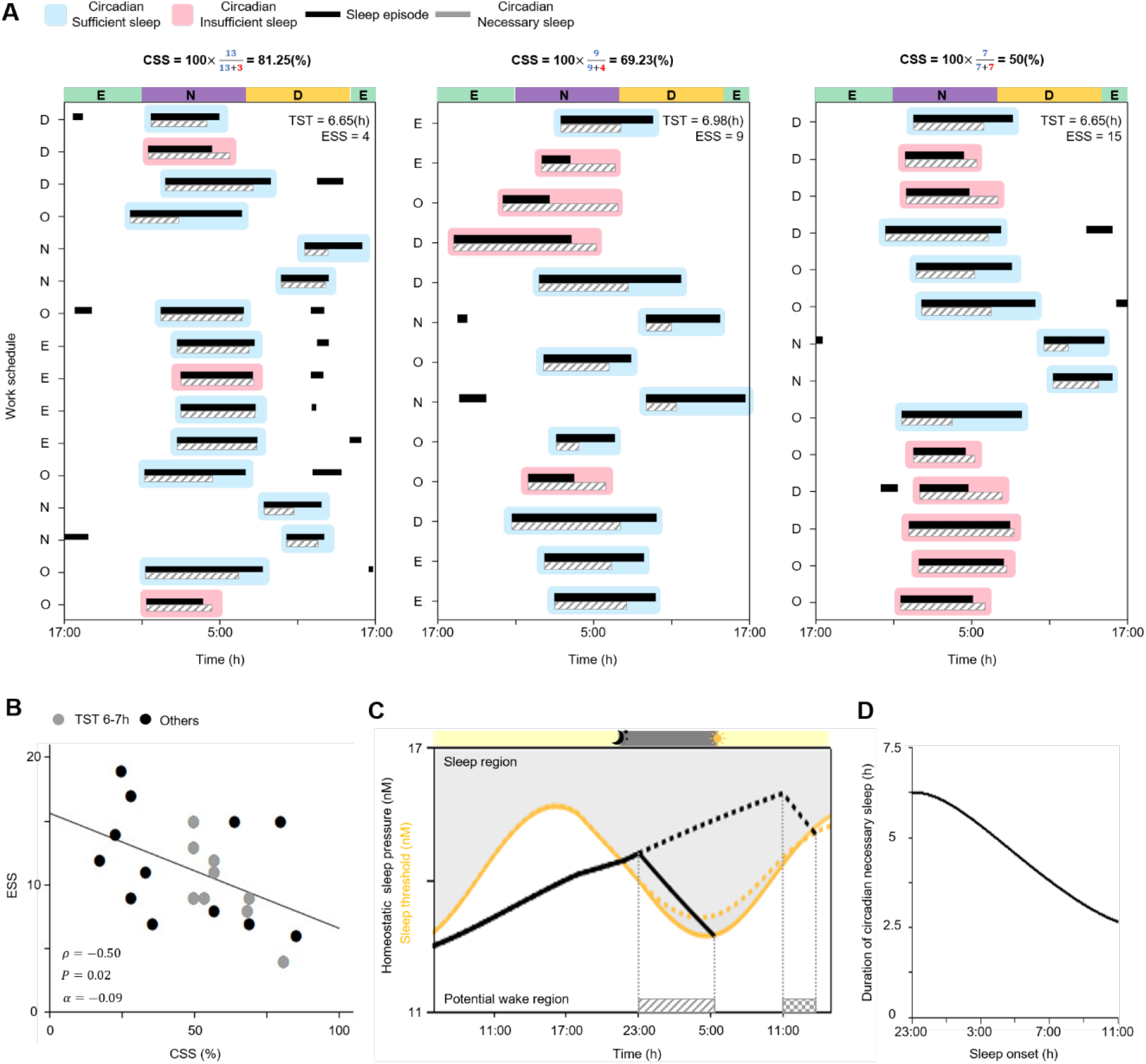
CSS is significantly correlated with the daytime sleepiness of shift workers. **(A)** Sleep-wake patterns of shift workers with similar TST but different ESS. Daily sleep episodes (black bars) whose duration is longer or shorter than the duration of circadian necessary sleep (gray striped bars) are categorized as either circadian sufficient sleep (blue shade) or circadian insufficient sleep (pink shade), respectively. As the fraction of circadian sufficient sleeps (i.e., CSS) decreased, daytime sleepiness (ESS) increased. D, E, N, and O denote the day shift (7:00-15:30 h), the evening shift (15:00-23:30 h), the night shift (23:00-7:30 h), and days off, respectively. **(B)** CSS had a strong and significant correlation with ESS. The line represents the least-square fitting line with the slope of *α. ρ* and *P* denote the Spearman’s rank correlation coefficient and *p* value of Spearman’s rank correlation test, respectively. **(C)** After regular 7-h sleep-wake patterns between 23:00 h and 6:00 h, simulations of sleep onset occurring at the usual time (23:00 h; solid line) compared to 12-h delayed sleep onset (11:00 h; dotted line). The duration of circadian necessary sleep needed after the delayed sleep (patterned bar) is much shorter than the duration of circadian necessary sleep needed after the regular sleep (striped bar). **(D)** As sleep onset is delayed from 23:00 h to 11:00 h, the duration of the predicted circadian necessary sleep gradually decreases by ~3.6 h.

To investigate this further in all the data recorded from the SMC participants, we developed a new sleep parameter, circadian sleep sufficiency (CSS), defined as the fraction of circadian sufficient sleeps in total sleep days during the study period. Indeed, over all the individuals, although there was some variation, CSS was significantly correlated with daytime sleepiness (*P* = 0.02; Figure 3B) unlike the other standard sleep parameters (Figure 1B–1D and Figure S1A-S1D). To the best of our knowledge, CSS is the first statistically significant sleep parameter for daytime sleepiness. Furthermore, CSS had a higher correlation with daytime sleepiness (*ρ* = −0.50) than any other standard sleep parameter previously discussed (Figure 1B–1D and Figure S1A-S1D). In particular, in 9 participants, despite having similar TST (6-7 h), as CSS increased, daytime sleepiness decreased (gray dots; Figure 3B).

### Sleep-wake patterns aligned with the circadian rhythm increase circadian sleep sufficiency

The duration of predicted circadian necessary sleep changed dramatically depending on the sleep onset time and previous sleep history (gray striped bars; Figure 3A). Thus, we further investigated how circadian necessary sleep was determined so that we could identify the sleep-wake patterns increasing CSS and thus decreasing daytime sleepiness. After regular 7-h sleep-wake patterns between 23:00 h and 06:00 h, we considered sleep onset occurring at the usual time (23:00 h; solid line; Figure 3C) compared to sleep onset delayed by 12-h (11:00 h; dotted line; Figure 3C). Unexpectedly, despite a dramatically increased homeostatic sleep pressure, the mathematical model predicted that the duration of circadian necessary sleep needed after the delayed sleep onset (patterned bar; Figure 3C) is much shorter than the duration of circadian necessary sleep needed after regular sleep onset (striped bar; Figure 3C). This shorter duration of circadian necessary sleep was due to the higher level of the sleep threshold during the day compared to the night, which is determined by the circadian rhythm (yellow lines; Figure 3C). That is, during the day, even after a short sleep, the homeostatic sleep pressure drops lower than the sleep threshold, and thus one can wake up without effort. Indeed, as sleep onset was delayed from 23:00 h to 11:00 h (Figure 3D), the duration of the predicted circadian necessary sleep gradually decreased by ~3.6 h. This indicates that the circadian rhythm is the key determinant of circadian necessary sleep. The importance of circadian rhythmicity has also been shown in previous studies reporting a decrease in sleep duration after sleep deprivation (Åkerstedt and Gillberg, 1981; Daan et al., 1984; Phillips and Robinson, 2008).

As circadian necessary sleep was mainly determined by the circadian rhythm, we hypothesized that sleep-wake patterns aligned with the circadian rhythm increase CSS and thus reduce daytime sleepiness. To investigate this, we simulated two different 3-day sleep-wake patterns. One follows three 6-h sleep episodes across three days regardless of sleep onset time, which is referred to as fixed sleep (Figure 4A). In the other simulation, sleep duration was adjusted according to the circadian phase of sleep onset, which is referred to as circadian sleep (Figure 4B). Despite having the same average sleep duration and sleep onset times, in the fixed sleep simulation only one sleep episode was categorized as circadian sufficient sleep (Figure 4A), while in the circadian sleep simulation all sleep episodes were circadian sufficient sleeps (Figure 4B). As a result, for the two circadian insufficient sleeps in the fixed sleep simulation, an awakening occurred before the homeostatic sleep pressure decreased below the sleep threshold (i.e., forced wakefulness). In real life, this situation can occur for example when using an alarm clock, or be caused by a disease, or stress (Foster, 2020; James et al., 2017; Skeldon et al., 2017; Van Dongen, 2004). Thus, after awakening from a circadian insufficient sleep, it takes some time for the individual to reach their potential wake region, where awakening would have occurred without effort (patterned bars; Figure 4A), and thus they may feel increased daytime sleepiness. In contrast, an individual awakening from circadian sufficient sleep is already in their potential wake region (patterned bars; Fig. 4A). Furthermore, after circadian sufficient sleep, the homeostatic sleep pressure was lower than after circadian insufficient sleep, and thus these individuals could be awake for longer before reaching their sleep threshold (e.g. ~32 min is longer before the third sleep; Figure 4A and 4B). Thus, the time awake in the potential wake region was ~8 h longer in the circadian sleep simulation compared to the fixed sleep simulation despite having the same average sleep duration (Figure 4C). This indicates that when the sleep-wake pattern is aligned with the circadian rhythm, the actual wake time is more likely to be aligned with the time of the potential wake region, and the duration of the potential wake region increases. As a result, there is less need for forced wakefulness, which may reflect daytime sleepiness.

**Figure 4.**
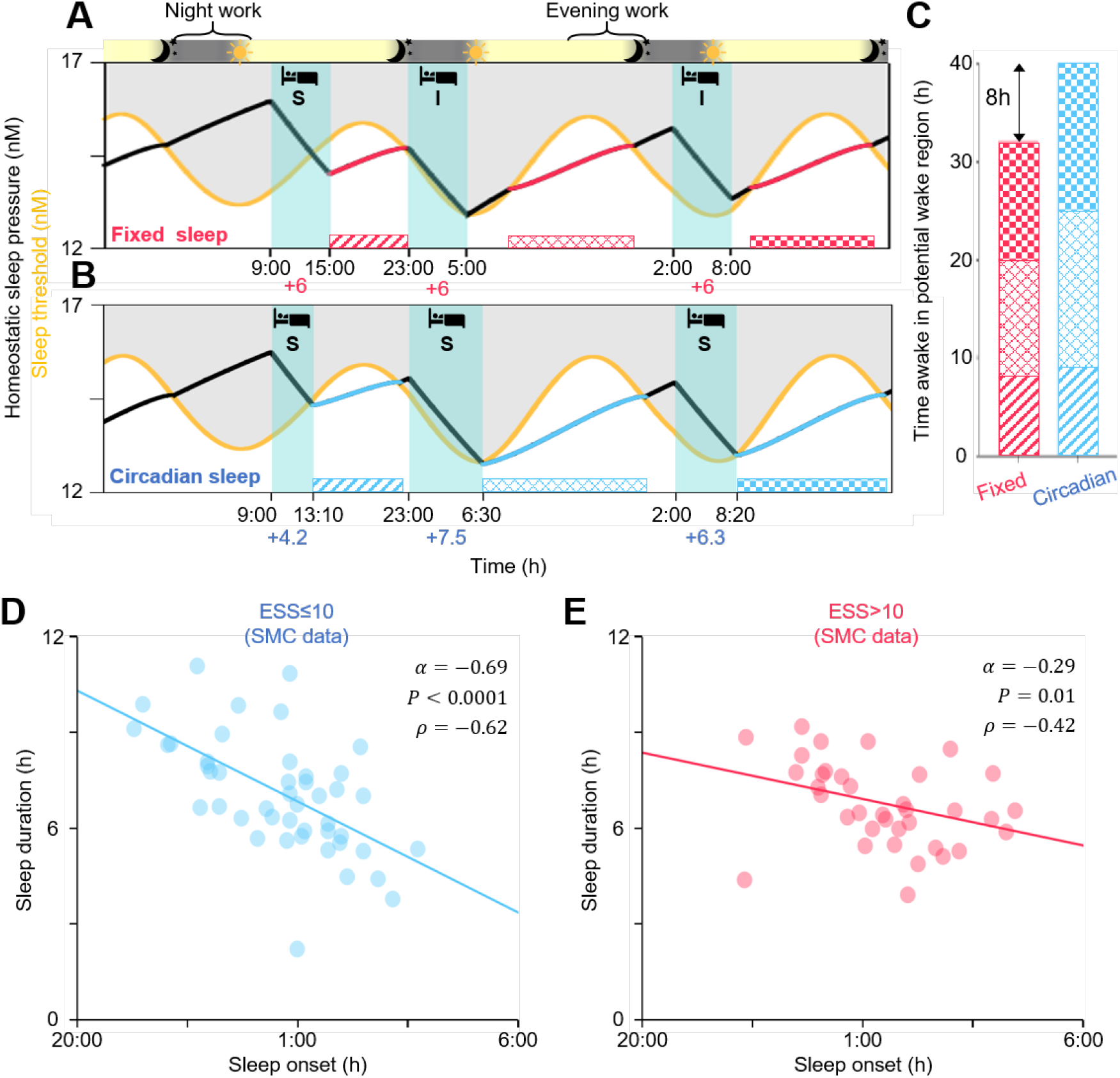
Sleep-wake patterns aligned with the circadian rhythm increase CSS and reduce daytime sleepiness. **(A** and **B)** Model simulations of three 6-h sleep episodes across three days regardless of sleep onset time (fixed sleep; A) and three sleep episodes with durations adjusted according to the circadian phase of sleep onset (circadian sleep; B). Although these two sleep-wake patterns have the same TST, two circadian insufficient sleeps (denoted as I) occur with the fixed sleep (A), but only circadian sufficient sleeps (denoted as S) occur with the circadian sleep (B). As a result, time awake in the potential wake region (patterned bars) is longer in the circadian sleep simulation (B) than in the fixed sleep simulation (A). **(C)** Quantification of the time awake in the potential wake region. **(D** and **E)** Alignment of sleep-wake patterns with the circadian rhythm of shift workers having similar TST (6-7 h) from SMC data (D and E; *n* = 9). The group without EDS (ESS≤10; blue dots; *n* = 5) show a much stronger negative dependence of sleep duration on the sleep onset time, compared to the group with EDS (ESS>10; red dots; *n* = 4). The number of analyzed main sleep episodes which were the longest sleep episodes of each day (noon-to-noon) were 45 (D) and 36 (E), respectively. The line represents the least-square fitting line with the slope of *α. ρ* and *P* denote the Spearman’s rank correlation coefficient and *p* value of Spearman’s rank correlation test, respectively.

### Sleep-wake patterns aligned with the circadian rhythm reduce daytime sleepiness

To investigate whether the better alignment of sleep with the circadian rhythm was associated with reduced daytime sleepiness, we analyzed the sleep-wake patterns of the shift workers from SMC. Specifically, we investigated whether a negative relationship between sleep onset time and sleep duration, as predicted by the mathematical model (Figure 3D), was stronger in the group without EDS (ESS≤10) compared to the group with EDS (ESS>10). For this comparison, we considered data only from shift workers having similar TST (6-7 h). Furthermore, sleep episodes before a day shift (7:00-15:30 h) whose wake onsets were usually triggered by an alarm clock, were excluded in this analysis to focus on the dependence of sleep duration on the circadian rhythm rather than forced sleep restriction following previous studies (Åkerstedt and Gillberg, 1981). As predicted, in the group without EDS, when sleep onset was delayed, which occurs often in shift workers, sleep duration clearly decreased following the circadian rhythm (*α* = −0.69; Figure 4D). This relationship was weaker in the group with EDS (*α* = −0.29; Figure 4E). This indicates that shift workers who aligned their sleep duration with their circadian rhythm had reduced daytime sleepiness (Figure 5). This provides personalized and flexible sleep-wake schedules reducing daytime sleepiness (Figure 5).

**Figure 5.**
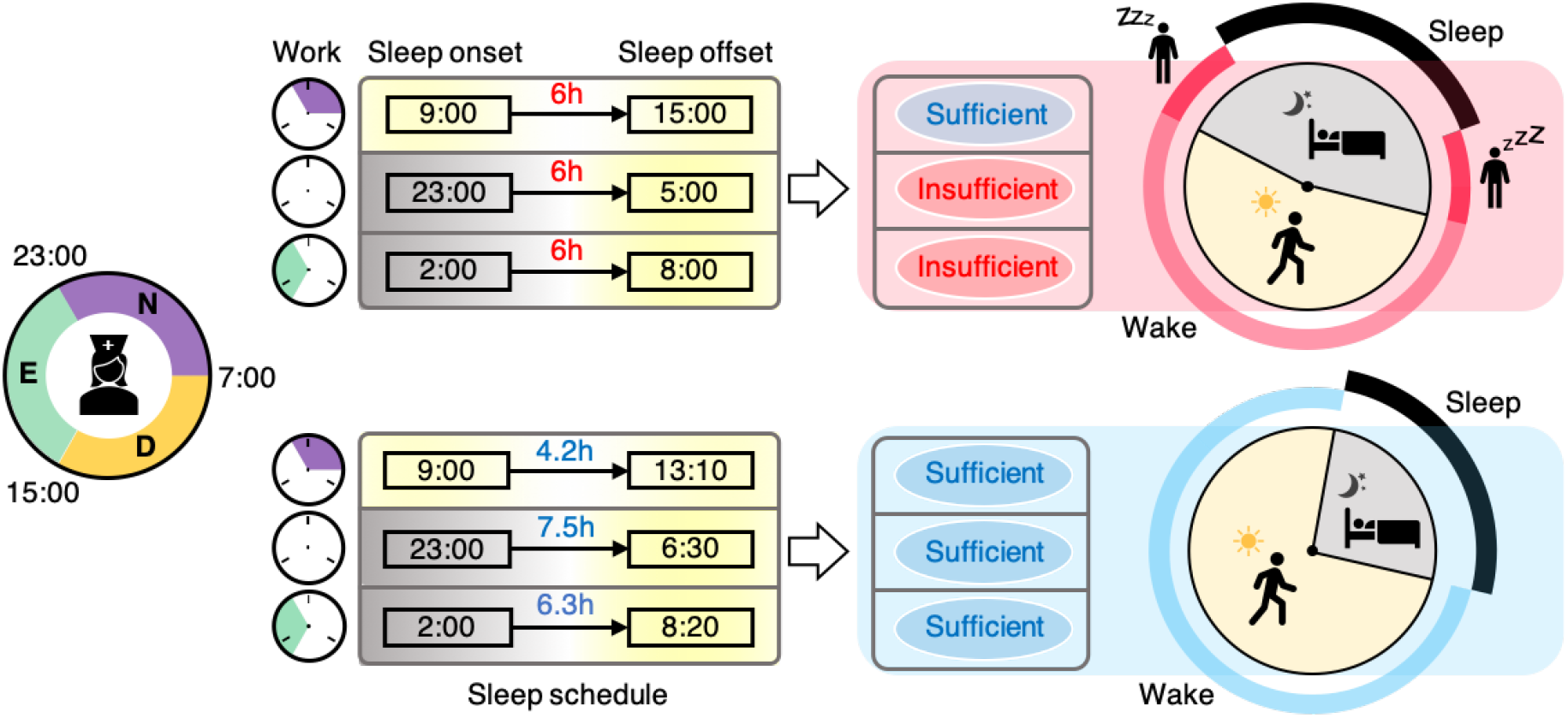
A sleep-wake pattern leading to circadian sufficient sleep reduces daytime sleepiness. Due to the alteration among day, evening and night shifts, sleep onset times of shift workers dramatically change. If they sleep for the same duration regardless of their sleep onset time, they frequently sleep less than the circadian necessary sleep, which is determined by their circadian rhythm and homeostatic sleep pressure, i.e., they have circadian insufficient sleep (top panels). Circadian insufficient sleep can be prevented if they actively change their sleep duration so their sleep-wake patterns match their natural sleep-wake cycle (bottom panels). As a result, they spend more time awake in the potential wake region when they feel less sleepy. In contrast, with circadian insufficient sleep, workers are awake in the sleep region, which requires wake effort and increases daytime sleepiness (top right). Note that the circadian insufficient sleep reduces the duration of the potential wake region as well.

## Discussion

We developed a user-friendly computational package that simulates the homeostatic sleep pressure and the circadian rhythm according to activity and light exposure, measured by wrist actigraphy (Figure 2 and Figure S6). Using this package, we found that shift workers whose sleep-wake patterns were aligned with their circadian rhythm had lower daytime sleepiness (Figures 3 and 4). Specifically, when they slept according to the computed duration of circadian necessary sleep, which was mainly determined by the circadian phase of bedtime, their sleep-wake patterns matched with their natural sleep-wake patterns (Figure 5). In this way, they awoke in the potential wake region when they would feel less sleepy and thus have lower daytime sleepiness (Figure 5). As these results were based on a retrospective study, it will be important to perform a prospective study investigating whether improving the alignment of sleep-wake patterns with the circadian rhythm reduces daytime sleepiness of individuals. The sleep-wake patterns aligned with the circadian rhythm were highly variable depending on various personal factors including average sleep duration, bed time, and environmental light exposure (e.g., Figure 3A). Importantly, our computational package can provide personalized and flexible sleep-wake schedules reducing daytime sleepiness.

With the computational package provided in this work, we were able to calculate the new sleep parameter CSS. As CSS quantifies the fraction of sleep episodes after which one can wake up without effort, it increases when sleep-wake patterns are better aligned with the circadian rhythm (Figures 4A and 4B). CSS showed a strong correlation with daytime sleepiness (Figure 3B) unlike standard sleep parameters such as TST and SL (Figure 1B–1C), indicating the importance of the circadian rhythm to understand daytime sleepiness, as has been emphasized in previous studies (Mairesse et al., 2014; Postnova et al., 2018; Puckeridge et al., 2011; Van Dongen, 2004). Importantly, the role of the circadian rhythm to understand complex aspects of sleep can be conveniently investigated with CSS. For instance, CSS can be used to investigate whether the circadian rhythm is a major source of inter-individual variations in sleep qualities and sleepiness depending on work schedules (Czeisler et al., 1982; Dunster et al., 2018; Vetter et al., 2015) and chronotypes (Vetter et al., 2015). Furthermore, the irregular sleep-wake patterns accompanied with the circadian misalignment have been considered as a major risk factor for insomnia, obesity, and cancer (James et al., 2017; Kecklund and Axelsson, 2016). How the risk of getting these diseases depends on sleep-wake patterns can also be effectively investigated with CSS.

Recent advances in wearable technology enable accurate real-time tracking of sleep-wake pattern and the circadian rhythm, which are critical components of our computational package (Cheng et al., 2021; Forger and Walch, 2020; Kim et al., 2020). A plethora of wearables have been developed to track sleep-wake patterns (Perez-Pozuelo et al., 2020). Recently, wearable devices measuring skin temperature and rest-activity successfully track the individual circadian rhythm during daily routine (Komarzynski et al., 2018). Even heart rate (Gao et al., 2014) and hormonal changes (Bariya et al., 2018), which are important factors for inferring the circadian rhythm, can also be tracked with wearables. The incorporation of these wearables and recently developed personalized sleep-wake mathematical models (Ramakrishnan et al., 2015) with our computational package can lead to the development of a smart alarm (Perez-Pozuelo et al., 2020). This will provide real-time personalized wake times, which align individual sleep-wake patterns with the circadian rhythm and thus reduce daytime sleepiness for those most suffering from it, including shift workers (Kato et al., 2012), patients of delayed sleep-wake phase disorder (Joo et al., 2017), Parkinson’s disease (Videnovic et al., 2017) or cancer (Sun et al., 2011).

### Limitations of the study

In this work, we developed the new sleep parameter CSS which has a significant correlation with daytime sleepiness of shift workers which was measured by ESS. Although ESS is one of the most widely used metrics to measure daytime sleepiness, it is a subjective metric. Future work will test whether CSS is still significantly correlated with daytime sleepiness measured with objective metrics such as psychomotor vigilance task or multiple sleep latency test. Additionally, when CSS is calculated, we did not consider the degree of circadian sleep sufficiency due to the relatively small size of data (e.g., 1h and 3h shorter sleep than circadian necessary sleep are considered as the same circadian insufficient sleep). It will be an important future work to identify a function describing the relationship between daytime sleepiness and the degree of circadian sleep sufficiency with large data.

### Resource Availability

#### Lead Contact

Further information and requests for resources and reagents should be directed to and will be fulfilled by the Lead Contact, Jae Kyoung Kim.

#### Materials Availability

This study did not generate new unique reagents.

#### Data and Code Availability

SMC data that support the findings of this study are available from the Lead Contact upon request. The MATLAB codes of the computational package are available in the following database: https://github.com/Mathbiomed/CSS. The GitHub repository will be made public when the manuscript is accepted.

## Methods

All methods can be found in the accompanying Transparent Methods supplemental file.

## Acknowledgments

This work was supported by the LG Yonam Foundation of Korea, the Human Frontiers Science Program Organization (RGY0063/2017), the National Science Foundation (DMS-1853506), and the Samsung Biomedical Research Institute grant (OTC1190671).

## Author Contributions

E.Y.J. and J.K.K. designed the study. S.J.C. and E.Y.J. collected and processed actiwatch data. J.H. and S.H.P. performed and H.H., V.B. and J.K.K. contributed to computational modeling and simulation. J.H. developed and H.H. contributed to the computational package. All authors analyzed the data. J.K.K. supervised the project. J.H. and J.K.K. wrote the draft of the manuscript, and all authors revised the manuscript.

